# Does arsenic contamination affect DNA methylation patterns in a wild bird population? An experimental approach

**DOI:** 10.1101/2020.12.08.415745

**Authors:** Veronika N. Laine, Mark Verschuuren, Kees van Oers, Silvia Espín, Pablo Sánchez-Virosta, Tapio Eeva, Suvi Ruuskanen

## Abstract

Pollutants, such as toxic metals, negatively influence organismal health and performance, even leading to population collapses. Studies in model organisms have shown that epigenetic marks, such as DNA methylation, can be modulated by various environmental factors, including pollutants, influencing gene expression and various organismal traits. Yet experimental data on the effects of pollution on DNA methylation from wild animal populations is largely lacking. We here experimentally investigated for the first time the effects of early-life exposure to environmentally relevant levels of a key pollutant, arsenic (As), on genome-wide DNA methylation in a wild bird population. We experimentally exposed nestlings of great tits (*Parus major*) to arsenic during their post-natal developmental period (3 to 14 days post-hatching) and compared their erythrocyte DNA methylation levels to those of respective controls. In contrast to predictions, we found no overall hypomethylation in the arsenic group. We found evidence for loci to be differentially methylated between the treatment groups, but for five CpG sites only. Three of the sites were located in gene bodies of zinc finger and BTB domain containing 47 (*ZBTB47*), HIVEP zinc finger 3 (*HIVEP3*) and insulin like growth factor 2 mRNA binding protein 1 (*IGF2BP1*). Further studies are needed to evaluate whether epigenetic dysregulation is a commonly observed phenomena in polluted populations, and what the consequences are for organism functioning and for population dynamics.

## Introduction

Environmental pollution can negatively affect organisms at multiple level of organization, from molecular and physiological level to performance, and even lead to population collapses (1-4). In wild populations, a largely unexplored mechanism mediating such pollution effects is the potential influence of the epigenome, such as DNA methylation. In human and animal models, the effects of pollution on the epigenome is studied extensively, and it has been discovered that methylation patterns can be changed by various environmental factors, including metal and organic pollutants and other early-life stressors (reviewed by 5-11). DNA methylation is the addition of a methyl (-CH3) group to the 5’ carbon site of cytosines catalyzed by DNA-methyltransferases, and is generally found to be negatively associated with gene expression (12). Variation in DNA methylation is linked to variation in phenotypes and behavior, and associated with the prevalence of various diseases, including cancers in humans and model animals (13-16). Epigenetic changes from early-life environment may persist and affect health throughout life-time and may even be transmitted to future generations (16), which could potentially contribute to explaining delayed or persistent effects of pollutants (e.g. 7). Yet the effects of pollutants on the epigenome have hardly been explored, and epigenetic research in wild animal populations is only emerging (8, 17-24).

Arsenic (As) is a global, persistent pollutant, distributed in the environment due to natural and anthropogenic sources such as mining, industrial activities or coal combustion (25) and the most highly ranked hazardous substances for animals and plants (26). Across organisms, arsenic can have negative consequences for basically all organ systems, often via causing oxidative stress, i.e. the imbalance between harmful reactive oxygen species (ROS) and antioxidant defenses, and cancer (27, 28).

Arsenic has been repeatedly observed to also modulate patterns of DNA methylation *in vitro* (e.g. 29) in laboratory animal models (with levels exceeding environmental levels, reviewed 30) and in studies on human populations (e.g. 31). Arsenic could influence DNA methylation via multiple pathways: (i) arsenic can change the DNA methylation of a cytosine via the depletion of the cellular availability of methyl groups. Biotransformation of arsenic to less toxic forms includes the addition of methyl group(s) (32) with the main methyl-donor for methylation of both arsenic and cytosines being s-adenosylmethionine (SAM). The high demand imposed on this molecule during the biotransformation process can then lead to a global DNA hypomethylation as shown in multiple (bio)medical studies in humans and mice (reviewed e.g. 5, 33). (ii) Arsenic could influence epigenetic signaling by targeting the zinc fingers of Tet proteins and perturbing the Tet-mediated oxidation of 5-mC (*in vitro*: 34-37). (iii) Furthermore, ROS created during arsenic biotransformation have been suggested to influence DNA methylation by creating aberrant modifications (humans: 38).

Pre/postnatal exposure to arsenic in humans is associated with epigenetic modifications related to early onset of diseases, which could have long-term consequences (reviewed in 39). For example, in humans, prenatal arsenic exposure led to global hypomethylation of inflammatory and tumor suppressor genes (40) and interfered with de novo methylation (41) in humans. Global hypomethylation can lead to chromosomal abnormalities, contributing to overall genomic instability and malignant transformations (reviewed in 32). Studies have demonstrated that widespread DNA hypomethylation induced by arsenic is also associated with promoter activation and involved in carcinogenesis (reviewed in 32). Arsenic-related hypomethylation of specific sets of genes has also be reported, and these include, for example, genes related to neural development (e.g. 42), mitochondria biogenesis (e.g. 43) and inflammation (e.g. 44). Despite the extensive data on model animals and humans, the potential effects of environmental arsenic on wild animals via epigenetic dysregulation has not been studied up to date.

We here investigated the effects of experimental early-life (post-natal) exposure to arsenic on genome-wide DNA methylation status in a wild population of great tits (*Parus major*). To our knowledge, this is the first study on the effect of arsenic on epigenetic marks in a wild population. We used a bird model, since birds have been successfully used in biomonitoring of pollution and its effects (e.g. 45). Arsenic exposure has been reported to negatively affect multiple fitness-related traits (growth, physiology, behavior and even egg-laying) in several bird species (reviewed in 28). For great tits specifically, we have previously reported (results from the current experiment) that in nestlings, arsenic exposure increased mortality, reduced wing growth (46) and decreased an intracellular antioxidant, catalase (47), but did not largely influence body mass, plasma biochemistry (vitamins) or other biomarkers of oxidative stress (46, 47). More specifically, we here experimentally exposed nestlings in non-polluted sites to environmentally relevant levels (1 µg/g body mass) of dietary arsenic during the entire post-hatching growth period, and compared their DNA methylation levels to respective controls. We used reduced representation bisulfite sequencing (RRBS) to assess genome-wide methylation and characterized differential methylation across CpG sites between the experimental and the control group. We predict that arsenic exposure will lead to genome-wide hypomethylation, and potentially specifically on gene/hubs related to development.

## Methods

### Arsenic treatment protocol and sampling

The study was conducted in the breeding season of 2015 (laying dates 4^th^ May –10^th^ June) in a nest box population of great tits (*Parus major)* in western Finland. Great tit is a small passerine bird and a popular model species in ecological and evolutionary research. Importantly, it is one of the few non-domesticated bird species, for which the genome and methylome are available (18, 48-49).

The arsenic exposure, dosages and sampling are described in detail in (46, 47). In short, the experiment was conducted in a nest-box population with known history of relatively low pollution levels (50). There are no air pollution samplers at the study sites but metal biomonitoring studies have been done in this area, for example measuring forest floor moss metal levels (a proxy for atmospheric fallout). In general, metal levels are relatively low in moss samples (e.g. for arsenic <0.5 μg/g in 2014; 51) while this value is exceeded in large areas in Central Europe (52). Mean topsoil arsenic concentration in the study site was 0.76 μg/g in 2014 (53).

Breeding was monitored, and from day 3 after hatching until day 13 whole broods were subjected to daily oral dosing with the following treatments: arsenic treatment (1 µg arsenic/g body mass in distilled water, N = 16 broods) or control treatment (distilled water, N = 16 broods). Dosing volumes were adjusted to estimated nestling mass based on average body masses at different ages from large dataset on long-term averages from the study population (54). Mass of individual nestlings was not measured every day to reduce handling time and disturbance to the nest. The volumes dosed to the controls were exactly the same as for treatments. We dosed the solution directly to the beak of the nestlings with a pipette. The range of volumes was 50–170 µl and did not exceed the recommended volumes (20 ml/kg, e.g. 55). The dose aimed to represent environmentally relevant exposure levels occurring in polluted areas in Europe: It was estimated combining data from several sources, such as (i) the lowest-observed-adverse-effect level for different effects on mammals (2-8µg/kg/day, 56), (ii) fecal arsenic levels reported for great tits at some metal polluted sites (reviewed in 28): In previous data, summarized in (28, Table 1), arsenic concentrations in feces of passerines are within the range of 0.1–1.4 ppm in unpolluted sites and 5–16 ppm in polluted areas. The levels measured in the samples from our experiment (ca 6.5 ppm, see results) overlap with these levels, suggesting that the treatment levels were environmentally relevant, at the lower end of the range. Yet, Sánchez-Virosta et al. (46) and Janssens et al. (57) report that great tit nestlings from polluted areas in Harjavalta and Belgium have arsenic levels up to 48-52 ppm, thus levels even this high are environmentally relevant. Other data sources were (iii) arsenic concentrations of food items (moth larvae, spiders and beetles) collected directly from parent great tits feeding their nestlings in the polluted area (46, 47), and (iv) a pilot experiment, to ensure that the levels were environmentally relevant and were not causing excessive mortality (46). Fecal matter was sampled 8 days after hatching for metal analyses (see below). DNA methylation was analyzed from red blood cells (RBCs, 14 d after hatching) to avoid sacrificing the individuals. Absolute methylation values between e.g. blood and liver or kidney and brain are highly correlated (48, 58, 59), just like changes in methylation in red blood cells and liver are correlated (59) and thus blood can be used as a proxy. Ten samples from the arsenic and ten from control treatment were selected for the DNA methylation analyses. These included five females and five males from each treatment (molecularly sexed, following 60). Only one nestling per nest was selected to avoid pseudoreplication. We made use of the knowledge on the fecal arsenic levels (see below), and selected individuals from 10 broods with highest arsenic concentrations from the arsenic treatment and 10 lowest concentrations from the control. All the dead nestlings found in the nests were collected and frozen at -20°C until necropsies could be performed in July 2015. Carcasses were necropsied to measure arsenic and metal concentrations in liver and bone in order to compare arsenic accumulation among groups and its distribution among tissues (46). The experiment was conducted under licenses from the Animal Experiment Committee of the State Provincial Office of Southern Finland (license number ESAVI/11579/04.10.07/2014) and the Centre for Economic Development, Transport and the Environment, ELY Centre Southwest Finland (license number VARELY/593/2015).

**Table 1.**
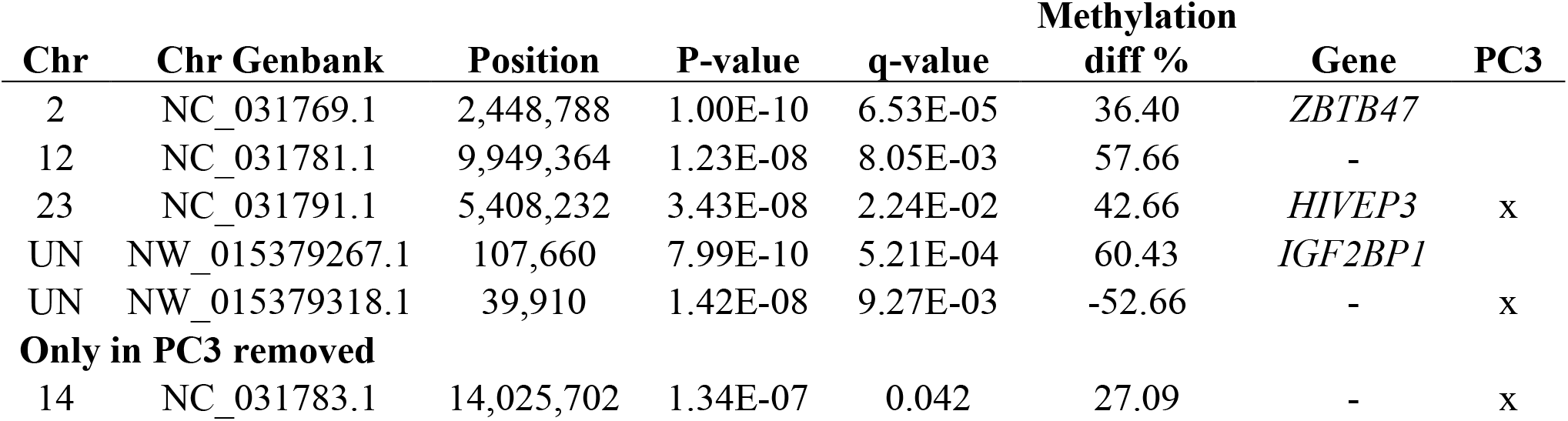
The differentially methylated CpG sites between arsenic exposed and control individuals. Methylation diff% refers to the methylation difference, comparing arsenic exposed to control group. Positive values therefore indicate hypermethylation in the arsenic treatment group compared to the control group. PC3 indicates sites that were significant after PC3 removal.

### Metal analyses

For detailed analyses, see Sánchez-Virosta et al. (46). Briefly, in both experimental groups, several fecal samples (any sex) from the same brood were combined to assess brood level metal exposure (total N = 32 broods). We determined the concentrations of arsenic, but also other metals to confirm that the levels of other metals were low and similar across the treatment groups (see 46). The determination of pollutants was conducted with inductively coupled plasma optical emission spectrometry (ICP-OES) with detection limit of 1 ppt (ng/l) and below. Calibration standards and certified reference materials were used for method validation. The levels of other measured metals (aluminium, lead, nickel, zinc, manganese, iron, copper) were low, and did not differ among the treatment groups (all t <0.88, all p <0.38).

### DNA isolation

DNA isolation was performed at the Center of Evolutionary Applications (University of Turku, Finland). We used RBCs given that previous studies suggest that blood shows similar methylation patterns as brain tissue in the study species (e.g. 80% similarity between brain and blood methylation in CpGs; 48, 49). DNA was extracted from 10-20 μl RBCs using the salt extraction method modified from (61). Extracted DNA was treated with RNase-I according to the manufacturer’s protocol. DNA concentration was measured fluorometrically with a Qubit High Sensitivity kit (ThermoFisher Scientific) and we assessed DNA integrity by running each DNA sample on an agarose gel.

### RRBS library preparation

We used a reduced representation bisulfite sequencing (RRBS) approach, which enriches for regions of the genome that have a high CpG content. We chose the RRBS approach because with the use of *MspI* as restriction enzyme, the method targets regions that are enriched for CpG sites. These regions are typically situated in or near the promotor regions, which has the advantage that CpGs in a relatively large proportion of the genes are covered (22, 62) making this a cost-effective method for detecting sites that are likely functional (16). It was previously shown in the study species that a vast majority of methylated Cs (97%) were derived from CpG sites in blood (48). Sequencing was conducted at the Finnish Microarray and Sequencing Center in Turku, Finland. The library preparation was started from 200 ng of genomic DNA and was carried out according to a protocol adapted from (63). The first step in the workflow involved the fragmentation of genomic DNA with MspI where the cutting pattern of the enzyme (C^CGG) was used to systematically digest DNA to enrich for CpG dinucleotides. After a fragmentation step a single reaction was carried out to end repair and A-tail (required for the adapter ligation) the MspI digested fragments using Klenow fragment (3’ => 5’ exo) following the purification of A-tailed DNA with bead SPRI clean-up method (AMPure magnetic beads). A unique Illumina TruSeq indexing adapter was then ligated to each sample during adapter ligation step to be able to identify pooled samples of one flow cell lane. To reduce the occurrence of adapter dimers, a lower concentration of adapters (1:10 dilution) was used than recommended by the manufacturer. These ligated DNA fragments were purified with bead SPRI clean-up method before putting samples through bisulfite conversion to achieve C-to-U conversion of unmethylated cytosines, whereas methylated cytosines remain intact. Bisulfite conversion and sample purification were done according to Invitrogen MethylCode Bisulfite Conversion Kit. Aliquots of converted DNA were amplified by PCR (16 cycles) with Taq/Pfu Turbo Cx Polymerase, a proofreading PCR enzyme that does not stall when it encounters uracil, the product of the bisulfite reaction, in the template. PCR-amplified RRBS libraries were purified using two subsequent rounds of SPRI beadclean-ups to minimize primer dimers in the final libraries. The high quality of the libraries was confirmed with Advanced Analytical Fragment Analyzer and the concentrations of the libraries were quantified with Qubit® Fluorometric Quantitation, Life Technologies. We used an average fragment size of 250-350 bp for sequencing.

### Sequencing

The samples were normalized and pooled for the automated cluster preparation which was carried out with Illumina cBot station. The 20 libraries were combined in two pools, 10 samples in each pool (treatments and sexes equally distributed between the pools) and sequenced in two lanes. The samples were sequenced with an Illumina HiSeq 2500 instrument using TruSeq v3 sequencing chemistry. Paired-end sequencing with 2 x 100 bp read length was used with 6 bp index run.

### Sequence data processing and differential methylation expression analysis

All the reads were checked for quality using FastQC (Babraham Bioinformatics) with multiQC (64), and low-quality sequences were trimmed with Trim Galore v. 0.4.4 (Brabraham Bioinformatics) by using --quality 20 --paired --rrbs settings.

The trimmed reads were mapped to the *Parus major* reference genome build 1.1. (http://www.ncbi.nlm.nih.gov/assembly/GCF_001522545.2) using Bismark (65) with default parameters. Methylation calling was conducted with Bismark, first with default settings with paired-end mode and overlap removal (--p --no_overlap). After this first calling round, we observed a methylation bias for the samples by plotting the methylation proportion across each possible position in the read. Based on the plotting, the three and two first bases of R1 and R2 respectively of the 5’ prime end were omitted and the first base in the R2 3’ prime end was also omitted in the final methylation calling. Thereafter, Methylkit (66) implemented in R was used for filtering and differential methylation analysis. We discarded bases that had coverage below 10x. To avoid a possible PCR bias we also discarded bases that had more than 99.9th percentile of coverage in each sample. Before differential methylation analysis we merged read counts from reads covering both strands of a CpG dinucleotide and CpGs needed to be covered with at least 8 samples per group (control and treatment).

Samples were thereafter clustered based on the similarity of their overall methylation profile by (i) using the clustering method ward.D in Methylkit’s clusterSamples -function and (ii) using principal component analysis (PCA) with Methylkit’s PCASamples -function. We also checked for lane and sex effect by using Methylkit’s assocComp -function where it checks which principal components are statistically associated with the potential batch effects such as the used lane and sex of the individuals. For the former, no missing data is allowed, thus we created a separate data object where all the individuals needed to be covered.

For analyzing differential methylation of CpG sites between control and arsenic treatment we used the beta-binomial model from DSS package (67) which is also included in Methylkit (calculateDiffMethDSS - function). DSS calculates the differential methylation statistics using a beta-binomial model with parameter shrinkage. Bonferroni correction was applied to account for multiple testing with q-value of 0.05. Furthermore, we also did the “tiling window analysis” in Methylkit where methylation information is summarized over tiling windows which are then used the in DSS analysis. We used the default values, win.size=1000, step.size=1000, cov.bases = 10 for the tiling and ran DSS again for these regions.

## Results

### Arsenic exposure

As reported in Sánchez-Virosta et al. (46, 47), dietary arsenic treatment successfully increased arsenic load as fecal arsenic levels were on average 10 times higher in arsenic exposure compared to control group (average±SD ppm: control 0.51±0.50, arsenic exposure 4.92±4.57, t_15.4_ = -3.83, p = 0.0015). In the subsample of nests selected for RRBS, the values were 0.47±0.37 ppm for control nests and 6.50±5.10 ppm for arsenic treatment, respectively. Furthermore, increased levels were also found in internal tissues: the mean (±SD) arsenic concentrations in liver were 4.19 ± 5.92 µg/g, d.w. (N=21) for arsenic exposure and 0.058 ± 0.100 µg/g, d.w. (N= 16) for control group, and in the bone 3.37 ± 3.85 µg/g, d.w. and 0.074 ± 0.103 µg/g, d.w., respectively (see Table 2 in 46). The levels were statistically significantly higher in arsenic exposure group compared to control group (p <0.001).

### Sequencing and mapping

The total number of read pairs was 341 million (Supplementary Table 1), varying from 14 million to 20 million per individual. After QC filtering the final number of read pairs was 337 million (Supplementary Table 1). The RRBS individual sequencing data have been deposited in NCBI (Number will be added later). Mapping efficiency was on average 46.15% and on average 3.1 million cytosines were covered before 10x coverage and percentile filtering. After filtering, 1.3 million cytosines were identified in CpG context. When combining the Cs from both strands and restricting our data to at least 8 individuals per group to be covered, we ended up having 652 655 CpGs.

### Sample clustering and differential methylation

Both the ward.D and PCA clustering methods showed that sample 14 from the treatment group was an outlier in its methylation profile (Supplementary Figure 1). That particular sample also had a low number of reads and showed lower duplication levels (Supplementary Table 1) and we therefore decided to exclude this sample from further analysis. No lane effect was detected, but PC3 (explained 0.23% of the variance) was associated with sex after Bonferroni correction (Supplementary Table 2, Supplementary Figure 2), mostly driven by two samples, ctrl_3F and test_16F, since after removing these two female samples from the data, PC3 was not significant anymore. Furthermore, when removing the PC3 from the data, three CpG sites were significant in the differential methylation analysis done with DSS: two of them were the same as when including all the PCs (see below, Table 1). The three other significant sites found below were not covered by all individuals as required in this PC-removal analysis.

In the differential methylation analysis when including all the PCs, five CpG sites showed a significant difference in methylation level with a q-value below 0.05 and percent methylation difference larger than 10% (Table 1, Figure 1, Supplementary Table 3). Lambda estimation was close to 1 (λ = 0.747, SE 0.000136) (Supplementary Figure 3), suggesting no systematic biases (λ>1 indicates bias). Four of these sites were hypermethylated (higher methylation in the arsenic treatment group) and one was hypomethylated (higher methylation in the control group). Three of the sites were located in gene bodies, namely zinc finger and BTB domain containing 47 (*ZBTB47*), HIVEP zinc finger 3 (*HIVEP3*) and insulin like growth factor 2 mRNA binding protein 1 (*IGF2BP1*) based on NCBI *P. major* annotation report 102. None of the regions from the tiling windows analysis were differentially methylated between control and treatment samples.

**Figure 1.**
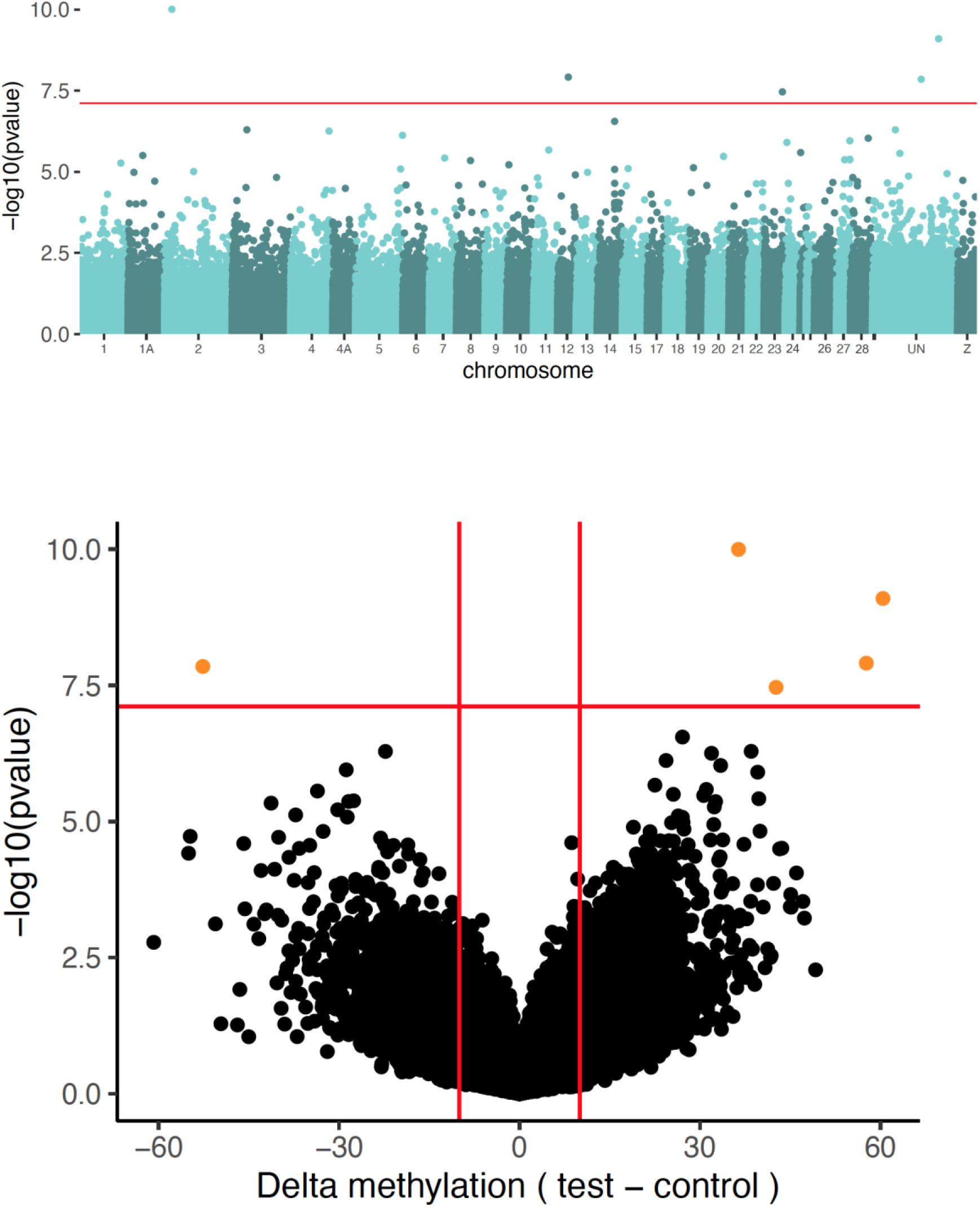
Plots of significance of CpG sites from the differential methylation analysis conducted with DSS implemented in Methylkit. (a) Manhattan plot with the significance of differential methylation of the arsenic treatment against the control pool, against the great tit reference genome version 1.1. The orange line depicts the genome-wide threshold based on a Bonferroni correction: 7.11. (b) A volcano plot of the significance against the absolute difference in methylation between the two pools, with delta methylation is arsenic treatment – control. Orange points are the genome-wide significant sites after Bonferroni correction and filtering for Delta methylation >10%.

## Discussion

We investigated whether early-life exposure to environmentally relevant levels of experimental arsenic affects DNA methylation in a wild vertebrate population. The experimental treatment increased arsenic levels significantly, but contrary to predictions, did not lead to overall hypomethylation. We found that treated individuals showed hypermethylation in four CpG sites and hypomethylation in one CpG site, indicating that increased levels of arsenic exposure appears to affect methylation at specific parts of the genome only. Yet also at these sites, the assumption of general hypomethylation was not met.

The lack of overall or site-specific hypomethylation may be explained by various factors: first, contrary to our predictions, the methyl donor s-adenosylmethionine needed for methylation may not have been limiting, potentially because oxidative status was not altered dramatically in all individuals. Indeed, as reported from the exact same experiment and samples by Sánchez-Virosta et al. (47), most biomarkers of oxidative status and damage in blood were only slightly (but not statistically significantly) elevated, and only the antioxidant enzyme catalase showed a significant decrease. In the future, sampling before and after exposure to e.g. pollutants may be advised to associate DNA methylation changes directly to changes in oxidative status, for example in adult birds (in contrast to developing animals where measurements are confounded by the changes in growth and associated changes in physiology).

Second, the response is likely to depend on the tissue type studied. For example, global hypomethylation in response to arsenic exposure is not consistently reported in blood: in humans, where blood leucocytes have been used to characterize arsenic associated changes no evidence for global hypo or hypermethylation was detected, yet arsenic was repeatedly reported to induce hypermethylation in various genes (especially promoters) (68), whereas global hypomethylation was detected in hepatic cells (69). Given that arsenic metabolism and SAM production mostly takes part in liver, we may expect tissue-dependent hypomethylation especially in liver, but not necessarily in other tissues. Unfortunately, we lack oxidative status measurements from the liver in this experiment. Studies have shown that absolute methylation values between e.g. blood and liver or kidney and brain are highly correlated (48, 58), just like changes in methylation in red blood cells and liver are correlated (59). Nevertheless tissue-specific methylation differences were larger for genes that are expressed in a tissue-specific way (48) and measuring methylation levels from red blood cells might therefore miss tissue-dependent genes whose expression is expected to change (59).

Furthermore, contrary to many previous studies in laboratory animals, this experiment was conducted with relatively low doses, mimicking exposure in polluted environments, whereas effects via SAM may only be apparent when levels are higher. Also, we were only interested in short-term, early-life effects while resident species inhabiting polluted environments during their whole life-span may show marked effects due to cumulative arsenic exposure. This is an interesting avenue for further research.

As the experiment was conducted in a wild population, in comparison to previous studies in laboratory, the environmental or genetic variability and potential variability across sexes may have masked some effects of the experimental treatments. Arsenic is known to have sex-dependent effects in many model systems (though predominantly in adult animals; reviewed e.g. 32). Furthermore, for example studies on mice report sex differences in DNA methylation patters in response to arsenic (e.g. 70), and general methylation differences among the sexes in young chickens (71). Yet, our initial models suggested that sex explained only a

very minor part of the variation in DNA methylation (and was therefore dropped from the final model), which suggests that in our data sex-bias is unlikely to strongly mask the effects. DNA methylation is known to be heavily influenced by the genetic background, for example in van Oers et al. (62), the majority of the variation between individuals was explained by genetic similarity. In the future, split-brood experimental designs may be used to distinguish genetic effects from environmental. The arsenic exposure applied (as measured from the fecal samples) was also at the lower range of variation if compared to polluted environments, which may contribute to the findings of only limited differences − yet mortality was increased with these levels, as reported in (46). We also report large variation in the fecal arsenic levels within the arsenic exposure treatment. Several factors may affect those levels, such as the time elapsed between last dosing and sampling, the times the nestling has been fed in that time and how many droppings they have produced, among others. Feces dropped soon after arsenic administration likely contain higher arsenic levels than later on.

We could annotate three of the five differentially methylated sites to genes. One of the genes, *IGF2BP1* is especially interesting as it is associated with development and growth: it has been showed that *IGF2BP1* plays important roles in various aspects of cell function, such as cell proliferation, differentiation, migration, morphology and metabolism (72, 73) but also embryogenesis and potentially even arsenic-related carcinogenesis (74, 75). *IGF2BP1* is abundantly expressed in fetal and neonatal tissues (73). Furthermore, two of the genes, *ZBTB47* and *HIVEP3* are both zinc-finger domains and are associated with transcriptional regulation (76). Epigenetic regulation of both *ZBTB47* and *HIVEP3* is known to be associated cancer (77, 78). Because our sample size in combination with a stringent correction for repeated sampling limits the power to detect subtle differences, we do expect to find a fraction of the number of differentially methylated CpGs (79).

All the three gene-related differentially methylated CpG sites were found in the gene body region, in both intron (*IGF2BP1*) and exons (*ZBTB47* and *HIVEP3*). Hypermethylation at CpG sites at promoter regions represses transcription of genes which is a well-known mechanism operating in many scenarios. DNA methylation at intergenic regions and gene bodies and its impact on gene expression is gaining more attention especially in cancer studies (80). Interestingly, a recent study on corals showed that gene body methylation was altered by environmental factors, which facilitated acclimatization and adaptation to different habitats (81). However, in great tits the DNA methylation observed in CpGs that are situated within gene bodies do not seem to affect gene expression (48), thus future studies are needed to determine the role of gene body methylation in gene expression control.

In conclusion, our study shows that early-life exposure to a toxic metal, arsenic, potentially affects fitness via DNA methylation changes in specific pathways, but not via an overall hypomethylation in the red blood cells. The effect might be more profound in other tissues that are more relevant to arsenic metabolism, such as liver. Thus, future studies should inspect other tissues as well. Other pathways of epigenetic alterations, known to be subject to arsenic-related alternations in vitro, such as histone acetylation (29) and micro-RNAs (82) could be further explored.

## ASSOCIATED CONTENT

**Figure S1**. Clustering of samples based on ward.D (a) and principal component analysis (PCA) in Methylkit

**Table S1**. Number of reads before and after read trimming with mapping and methylation calling success of Bismark. CpG site filtering was done with Methylkit.

**Table S2**. Results from the tests on the effects of sequencing lane and sex.

**Table S3**. A full description of all differentially methylated sites.

## Data accessibility

Data will be deposited in Dryad and in Genbank upon acceptance.

## Author contributions

SR, SE, PSV, VNL, MV, KvO and TE designed the study. TE, SE and PSV collected the data. SR and KvO designed the sequencing. VNL and MV conducted the bioinformatic analyses. SE and PSV conducted metal analyses. KvO and VNL provided the genome resources. VNL, MV, KvO and SR interpreted the data. VNL and SR wrote the first draft. All authors contributed to writing the manuscript.

## Acknowledgements

We thank Miia Rainio and Jorma Nurmi for their efforts in helping us with fieldwork and Fleur Gawehns-Bruning and William Sies for assistance with the methylation analysis. We also thank the Center for Evolutionary Applications for molecular work and Finnish Functional Genomics Centre for sequencing services. Our study was financed by KONE foundation (SR) and Academy of Finland (TE: project 265859). SE is currently funded by *Ministerio de Ciencia, Innovación y Universidades* (IJCI-2017-34653 to SE).

