## Supplementary material for "Does arsenic contamination affect DNA methylation patterns in a wild bird population? An experimental approach" (from another version of this preprint: version 1, 2020-12-09)

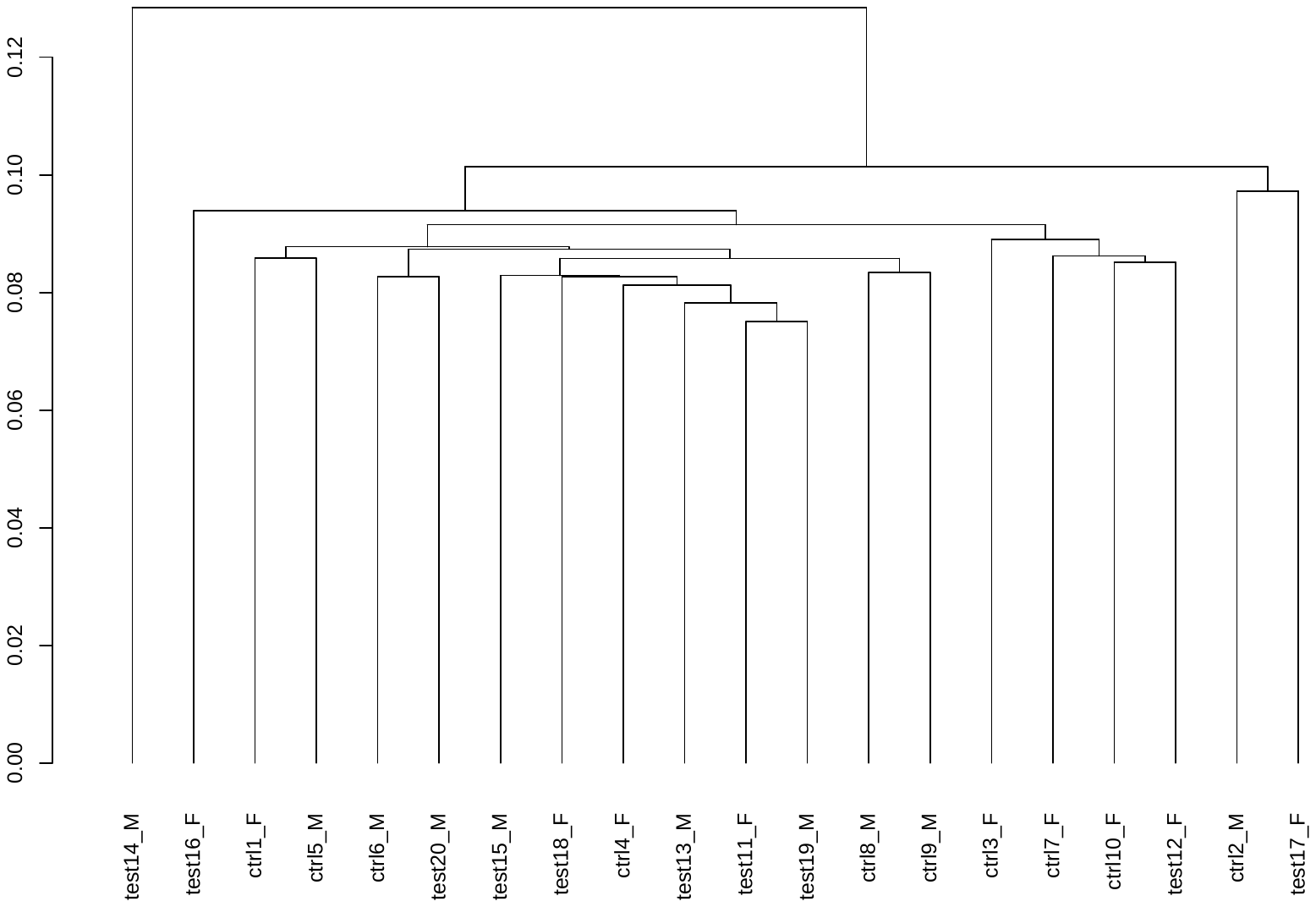

**B**

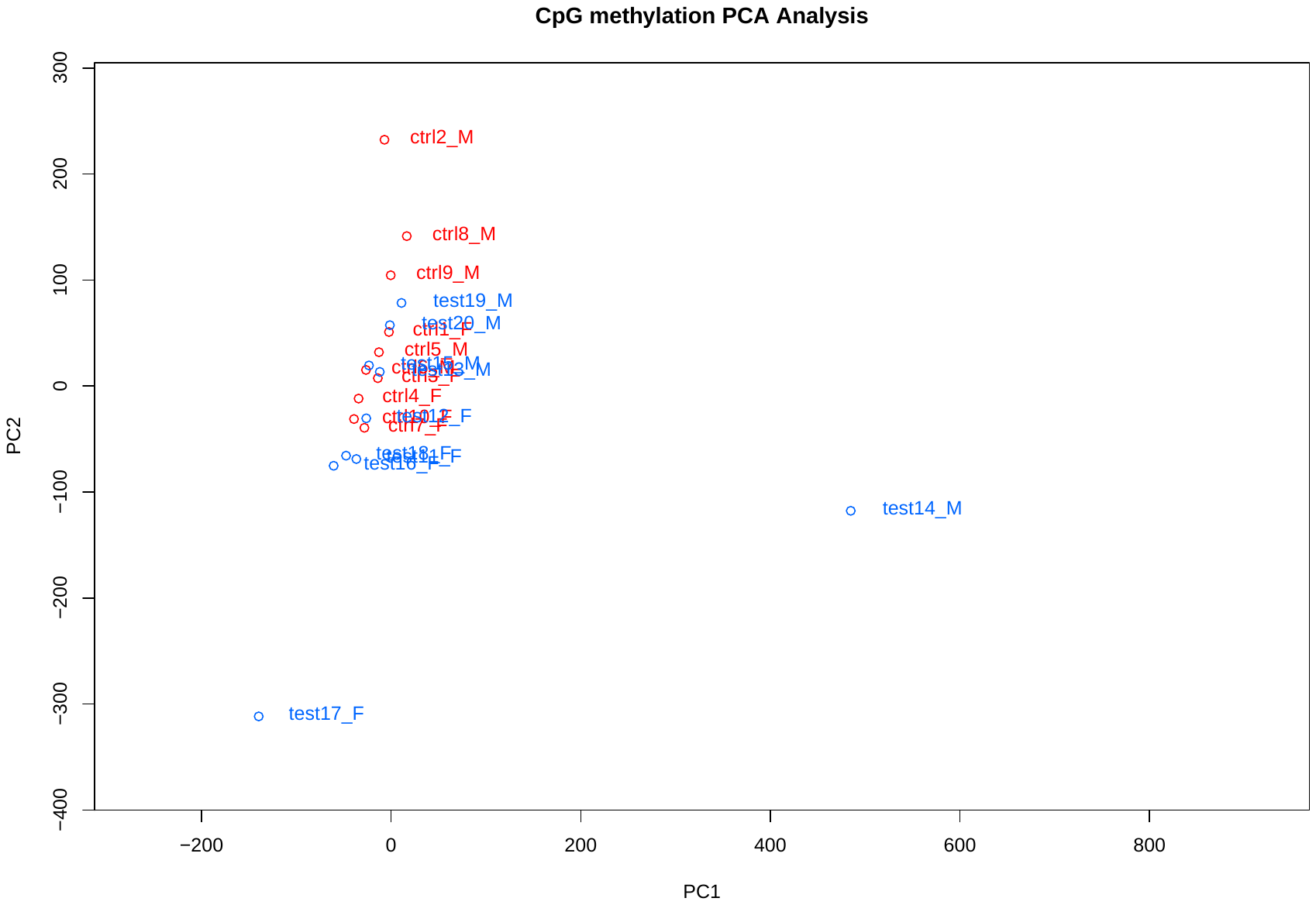

**Supplementary Figure 1.** Clustering of samples based on ward.D (a) and principal component analysis (PCA) (b) in Methylkit. Names of individuals represent treatment (test = treated individuals (blue) and ctrl = control individuals (red)), number and sexes (F = female and M = male). The PC1 explained 9% of the variance and that of PC2 is 7.6%.

**A**

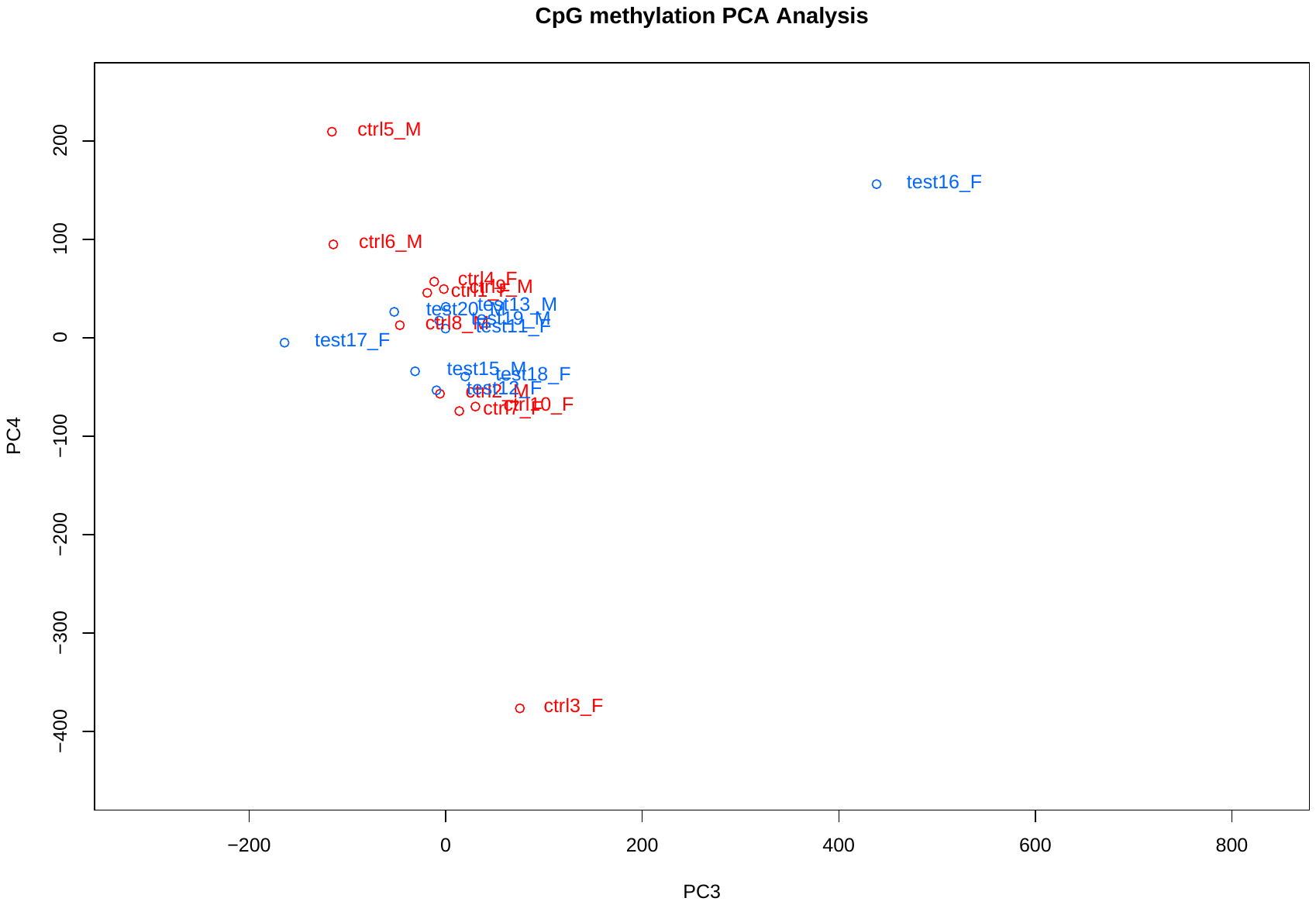

**Supplementary Figure 2.** Clustering of samples based on principal components PC3 and PC4 by PCA analysis in Methylkit after removing outlier sample 14. Names of individuals represent treatment (test = treated individuals (blue) and ctrl = control individuals (red)), nuumber and sexes (F = female and M = male). The PC3 explained 6.2% of the variance and that of PC4 is 5.9%.

**A**

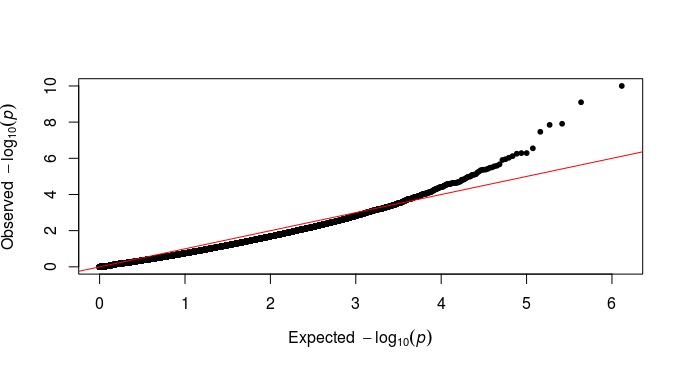

**Supplementary Figure 3.** QQ plot of observed vs. expected -log10 transformed P-values. Lambda = 0.747 (SE 0.00013)
